# Pan-genome Analysis Reveals Hidden Diversity and Selection Signatures of Auxin Response Factors (ARFs) Associated with Breeding in Barley

**DOI:** 10.1101/2025.08.12.669851

**Authors:** Kenan Tan, Zhenru Guo, Thorsten Schnurbusch

**Author notes:** These authors contributed equally to this work (Kenan Tan, Zhenru Guo).

## Abstract

Auxin Response Factors (ARFs) play a pivotal role in regulating plant growth and development, yet their evolutionary dynamics and functional divergence remain poorly understood in barley (*Hordeum vulgare* L.) In this study, we conducted a comprehensive genome-wide analysis of the ARF gene family across a 76-accession barley pan-genome. By integrating gene presence/absence variation (PAV) and copy number variation (CNV), phylogeny, expression profiling, transposable element (TE)-mediated regulation, and selection signatures, we characterized 1,911 ARF-coding genes and their structural, transcriptional, and functional variation at the population level. Phylogenetic analysis revealed lineage-specific expansion and dynamic duplications, particularly within the HvARF13 clade. Co-expression networks and tissue-resolved transcriptomes showed that many *HvARFs* are preferentially expressed in inflorescence and meristematic tissues. Selective sweep and haplotype analyses identified *HvARF3* as a candidate gene under selection during European barley breeding. A favorable haplotype of *HvARF3*, enriched in European cultivars, was significantly associated with increased grain size and weight, demonstrating the utility of pan-genome-enabled frameworks for accelerating gene-trait association and candidate gene discovery. This study highlights the power of multi-omics integration in decoding gene family complexity and provides valuable insights for functional genomics and trait improvement in cereal crops.

## Introduction

Auxin response factors (ARFs) are a family of plant-specific transcription factors that mediate auxin signaling by binding to auxin-responsive elements in gene promoters, thereby regulating a broad array of developmental and physiological processes. These include organogenesis, embryogenesis, apical dominance, root and shoot development, and reproductive differentiation (Cancé et al., 2022). Typically, ARF proteins harbor three functional domains: a conserved N-terminal B3 DNA-binding domain (DBD), a variable middle region that acts as either a transcriptional activator (glutamine-rich, Q-rich) or repressor (serine/threonine/proline-rich, STP-rich), and a C-terminal PB1 (Phox and Bem1) domain that facilitates homo- and hetero-dimerization with other ARFs or Aux/IAA proteins (Chandler, 2016). This domain architecture is remarkable conserved from early land plants to higher angiosperms, underscoring the fundamental roles ARFs play across plant evolution (Li et al., 2023).

In numerous species—such as *Arabidopsis thaliana*, rice (*Oryza sativa* L.), maize (*Zea mays* L.), and wheat (*Triticum aestivum* L.)—*ARF* genes have been systematically identified and functionally characterized (Gidhi et al., 2021; Man et al., 2025; Okushima et al., 2005; Wang et al., 2007; Xing et al., 2011). These studies revealed the involvement of specific ARF members in regulating key agronomic traits such as grain size, inflorescence architecture, and stress responses. For example, *OsARF3*, *OsARF4*, *OsARF6*, *ZmARF24*, *ZmARF2*, *ZmARF17*, and *TaARF25* have been reported to regulate grain/kernel traits (Gao et al., 2023; Gu et al., 2023; Hu et al., 2018; Jia et al., 2021; Qiao et al., 2021; Wang et al., 2024). Notably, *OsARF3* is also involved in balancing the trade-off between abiotic and biotic stress responses. *OsARF12*, *OsARF17*, *OsARF25*, and *OsARF10* have been shown to affect stigma development (Guo et al., 2022; Zhao et al., 2022). *ZmARF28* can pleiotropically modify plant architecture by single amino acid substitution, its mutant shows reduced plant height, increased leaf number and changed inflorescence identity (Prigge et al., 2025). These functional insights demonstrate the evolutionary versatility and their potential for agronomic improvement of *ARF* genes across cereals. Despite its agronomic and historical significance, as one of the first domesticated crops (Lister et al., 2018), barley (*Hordeum vulgare* L.) has not yet seen comprehensive systematic characterization of its *ARF* gene family. Although *ARF* genes have been implicated in regulating key yield and fitness traits in related cereals, their characterization in barley remains limited, highlighting a significant knowledge gap. Given the known roles of *ARF* genes in other species, elucidating the diversity and function of the ARF protein family in barley could reveal previously unknown regulators of important developmental processes and contribute directly to crop improvement efforts.

Recent advances in high-throughput sequencing and the development of pan-genome and pan-transcriptome resources have transformed plant genomics by enabling species-wide resolution of genetic diversity (Marks et al., 2021). Traditional single-reference-based analyses are insufficient to capture the full extent of gene family variation, as they often miss presence/absence variants (PAVs), copy number variations (CNVs), or population-specific alleles. Pan-genomic resources—such as those recently generated for barley—offer a more comprehensive view of structural and functional diversity at both the gene and genome levels (Jayakodi et al., 2024; Jayakodi et al., 2020). In parallel, transcriptome datasets across tissues, genotypes, and even at single-cell resolution now make it possible to explore gene expression divergence, regulation, and potential adaptation at an unprecedented scale.

In this study, we performed a comprehensive, population-scale analysis of the ARF protein family in barley using integrated multi-omics data. We systematically identified and annotated genes and proteins of the ARF family across the 76 barley pan-genomes, revealing approximately 30% more members than in previous studies. By constructing high-resolution phylogenies and characterizing structural variation, expression patterns, *cis*-regulatory elements, and selective signals, we explored the evolutionary and functional dynamics of ARFs in barley. Our findings revealed significant differences in natural selection pressures and genomic diversity across distinct barley populations. Notably, we identified a superior allele of *HvARF3* that appears to have undergone selection in European cultivars, with significant association to increased grain size and weight. These findings demonstrate how integrative pan-genome and transcriptome analyses can illuminate both conserved gene family evolution and novel, population-specific adaptations with agronomic relevance. Our results provide a comprehensive panoramic view of the ARF protein family and establish a foundation for future studies on auxin signaling, inflorescence development, and molecular breeding in barley.

## Results

### Identification and validation of *ARF* genes and their PAVs/CNVs across 76 barley pangenome assemblies

To comprehensively catalog *ARF*-coding genes in barley, we employed a domain-based gene discovery approach by integrating three major protein family databases—Pfam (PF02362, PF06507, PF02309), ProSiteProfiles (PS50863, PS51745), and PANTHER (PTHR31384) —using the recently released barley pan-genome v2 database (BPGv2) (Jayakodi et al., 2024). All candidate sequences were further validated against the PanBARLEX online resource (https://panbarlex.ipk-gatersleben.de/<u>#</u>) to ensure annotation accuracy. Given that the conventional reference genome (Morex) lacks two *ARFs* due to genotype-specific presence/absence variations (PAVs), we selected the European modern spring barley cultivar Barke for gene nomenclature. Barke harbors all identified *ARF*-coding genes and has been extensively characterized at the transcriptomic level (Coulter et al., 2022; Guo et al., 2025). To maintain cross-species consistency, gene names were assigned based on the wheat gene nomenclature system (Boden et al., 2023) and aligned with orthologous ARF genes in rice (Wang et al., 2007).

Compared to previous single-genome based analyses (Tombuloglu, 2019), our pan-genomic approach uncovered approximately 30% more *ARF* genes, underscoring the power of multi-genotype analysis to capture greater gene diversity. We retained 26 full-length *ARFs* containing at least one conserved ARF-related domain in ≥10% of accessions for downstream analysis. Based on their frequency across the 76 barley genomes, these genes were classified into core, soft-core, and shell categories following established dispensability thresholds (Marroni et al., 2014) (Figure 1A, E). Notably, approximately ∼80.7% of ARFs belonged to the core or soft-core categories, being present in 76 or 75 genomes, respectively. This high evolutionary conservation highlights the essential roles of ARFs for barley development and fitness.

**Fig. 1.**
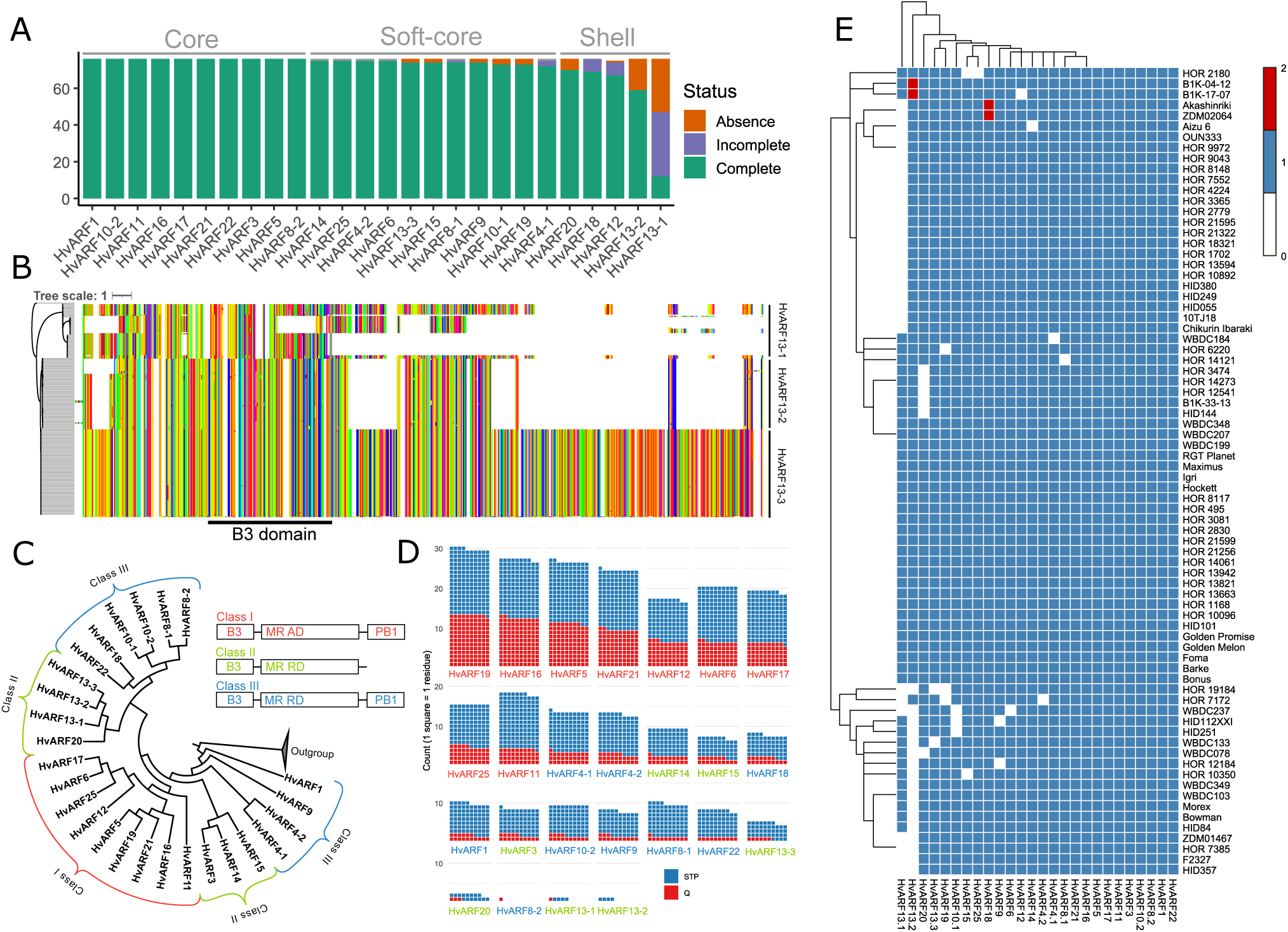
Identification and characterization of *ARF* genes in the barley pan-genome. (A) Distribution of PAVs for each *ARF* gene across 76 barley accessions. Genes were categorized as core (present in all 76 genomes), soft-core (present in ≥95% of genomes), or shell (present in 10%-95% of genomes). (B) Amino acid sequence alignment of HvARF13-1, HvARF13-2 and HvARF13-3. Different colors represent different amino acid residuals. (C) Phylogenetic tree of *ARF* genes in the cv. Barke, classified according to gene structure. (D) Comparison of Q residual counts versus S/T/P residuals in the middle regions of ARF proteins in cv. Barke. (E) Heatmap showing the presence or absence of each *ARF* gene across the 76 barley genotypes.

An exception was the HvARF13 clade, which exhibited extensive structural and presence variation. While *HvARF13-3* was highly conserved (found in 74 genotypes), *HvARF13-1* and *HvARF13-2* were present in only 47 and 59 accessions, respectively. Although HvARF13-1 and HvARF13-2 are shorter than HvARF13-3, all three retain an intact B3 DNA-binding domain in at least some genotypes and are expressed at transcriptome level, suggesting potential functionality (Figure 1B). This structural variation reflects dynamic gene duplication events and copy number variations (CNVs) within the barley ARF family. To validate these PAVs, we extracted 5-10 kb flanking genomic sequences and performed local alignment. A 479 bp deletion was identified at the 3′ end of *HvARF13-2* in several genotypes, including Morex (Figure S1B), likely explaining its absence in previous annotations. A larger deletion was detected for *HvARF13-1* in some accessions, leading to complete gene loss (Figure S1A). Additionally, a truncated gene (HORVU.BARKE.PROJ.7HG00820280) lacking conserved domains was likely misannotated and was excluded after sequence and domain verification.

To explore functional implications of ARF protein diversity, we examined the composition of the middle region in each Barke ARF protein, combining domain annotation with phylogenetic analysis using rice homologs (Figure 1C). We specifically quantified glutamine (Q) and serine/threonine/proline (STP) residue enrichment, which are indicative of functional differentiation (Chandler, 2016; Shen et al., 2010). ARFs with Q-rich middle regions were classified as Class I (typically transcriptional activators; e.g. *HvARF19*), whereas those enriched in STP residues were categorized as Class II or Class III (transcriptional repressors; e.g. *HvARF1*) (Figure 1D). In total, 26 well-characterized ARFs with high-confidence annotations were selected for our subsequent analyses.

### Phylogenetic and *Ka*/*Ks* analyses reveal evolutionary dynamics and selection pressures on ARF proteins in barley

To elucidate the evolutionary relationships within the ARF gene family in barley, we performed a comprehensive phylogenetic analysis based on full-length ARF protein sequences derived from 76 barley genotypes. Despite extensive sequence variation among accessions, all ARF proteins could be reliably assigned to their respective clades, consistent with ARF classifications established in other plant species (Wang et al., 2007) (Figure 2A). Comparative genome analyses across *Arabidopsis*, rice, and barley revealed substantial changes in ARF copy number, with both gene losses and expansions observed. Notably, several *Arabidopsis* ARFs, including *AtARF9*, *AtARF13*, *AtARF14*, *AtARF23*, *AtARF12*, *AtARF22*, *AtARF15*, *AtARF10*, *AtARF20*, and *AtARF21*, were completely absent in both rice and barley (Figure 2A), suggesting lineage-specific gene loss events following the divergence of grasses and dicots (Aravind et al., 2000; Finet et al., 2013). This pattern likely reflects evolutionary streamlining of the *ARF* repertoires in grasses and possibly the adoption of distinct auxin signaling strategies (McSteen, 2010).

**Fig. 2.**
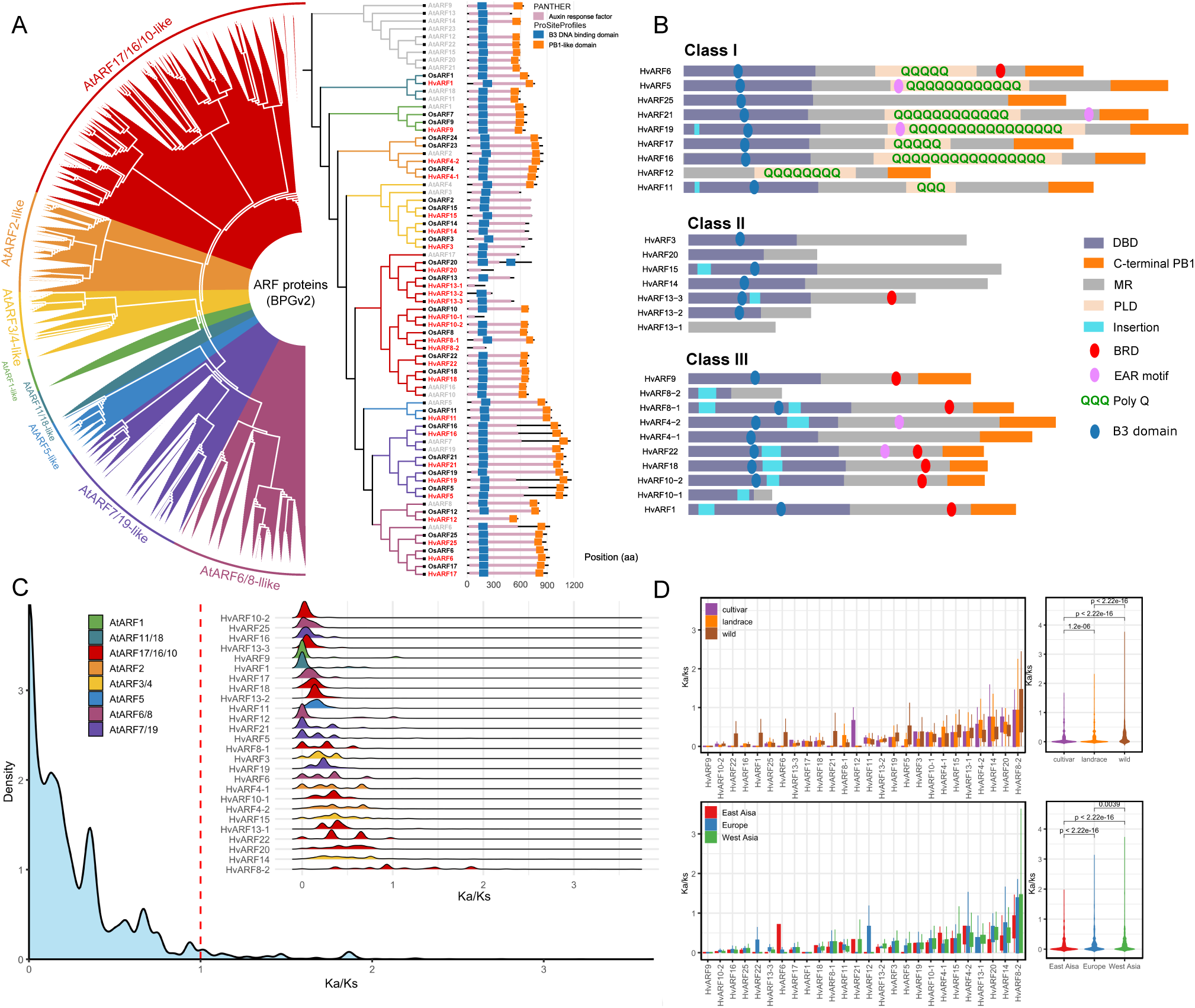
Phylogenetic relationships, protein structure and *Ka*/*Ks* analysis of ARF proteins. (A) Phylogenetic tree of ARFs identified from 76 barley genotypes (left) and a comparative tree including ARF proteins from barley (cv. Barke), rice and *Arabidopsis* (right). Conserved protein domains of ARF proteins from Barke, rice and *Arabidopsis* are shown to the right of the phylogenetic tree. (B) Schematic organization of ARF-MR domains in barley cv. Barke. DBD, DNA-binding domain containing the B3 domain and nearby dimerization domains; MR, middle region; PLD, prion-like domain; EAR motif, putative EAR motif characterized by the short sequence LxLxL/DLNxxP; BRD, B3 repression domain; B3 domain, the core sequence of DBD domain. (C) Distribution density of *Ka/Ks* values for each ARF protein and the overall *Ka/Ks* distribution across the ARF protein family. (D) *Ka/Ks* ratios of individual ARF proteins, including comparisons among cultivars, landraces and wild barley, as well as among geographic groups: East Asia, Europe and West Asia.

Interestingly, the AtARF5 clade retained a single-copy ortholog across all three species tested here, indicative of strong purifying selection and a critical, possibly irreplaceable, role in plant development. In contrast, barley showed expanded *ARF* families compared to rice, particularly in *HvARF4*, *HvARF8*, *HvARF10*, and *HvARF13*. Among these, *HvARF4-1* and *HvARF4-2* represent a rare example of full-length gene duplication, with both paralogs retaining all key functional domains, including the B3 DNA-binding domain and PB1-like domain (Figure 2A, 2B). Other duplicated genes, such as members of the HvARF13 clade, often exhibited partial structures lacking one or more conserved domains, suggesting potential subfunctionalization or neofunctionalization. Notable, most of the observed barley-specific ARF duplications clustered within the AtARF17/16/10-like clade, which includes *AtARF10*, *AtARF16*, *AtARF17*, and *OsARF18*—a group known to function as negative regulators of plant development and seed germination (Huang et al., 2016; Liu et al., 2007; Liu et al., 2013; Mallory et al., 2005). The increasing copy number of this clade from *Arabidopsis* to rice and barley suggests an adaptive expansion, potentially driven by environmental selection pressures or functional diversification. To investigate the variation among ARF protein in barley, all ARF protein sequences were analyzed and are presented in a schematic diagram (Figure 2B) (Cancé et al., 2022). The DBD domain is relatively conserved in class I ARFs, whereas almost all class III ARFs contain a short insertion within this region, suggesting greater structural stability and conservation in class I. Notably, HvARF25, although belonging to class I, has a relatively high content of glutamine (Q) residues but does not form a poly-Q structure. Most class III ARFs possess the B3 repression domain (BRD, motif pattern: R/KLFG) (Choi et al., 2018), a characteristic indicator of repressor function. However, HvARF4-1 and HvARF4-2 are exceptions: instead of the canonical R/KLFG motif, each contain a single amino acid substitution that produces a modified KIFG motif, potentially indicating a functional divergence.

To assess selection constraints acting on ARF proteins, we calculated the ratio of non-synonymous (*Ka*) to synonymous (*Ks*) substitution rates (*Ka/Ks*) across different barley populations. *Ka/Ks* values below, equal to, or above 1 are generally interpreted as indicative of purifying, neutral, or positive selections, respectively (Kimura, 1985). Overall, most ARF proteins exhibited *Ka/Ks* ratios well below 1, indicating strong purifying selection acting to conserve gene function across genotypes (Figure 2C). This evolutionary constraint underscores the essential regulatory roles of ARF proteins during barley development and adaptation. However, *HvARF8-2* was a notable exception, with *Ka/Ks* ratios frequently exceeding 1. The lack of a B3 domain in this gene may have relaxed functional constraints, permitting rapid evolution under natural or selective breeding. This gene may represent an example of lineage-specific divergence or a recently neofunctionalized paralog. Further comparison of *Ka/Ks* ratios among wild barley, landraces, and modern cultivars revealed a statistically significant decline from wild accessions to cultivars, indicative of intensified purifying selection during domestication and breeding. A similar trend was observed across geographic regions: accessions from East Asia exhibited lower *Ka/Ks* values than those from European and West Asia (Figure 2D), suggesting differential selection pressures linked to regional adaptation.

Interestingly, *HvARF12* deviated from this general pattern, showing elevated *Ka/Ks* ratios in cultivars compared to wild or landrace accessions. Its rice ortholog, *OsARF12*, is functionally redundant with *OsARF17* and *OsARF25*, and has been implicated in auxin-mediated regulation of tiller angle, as well as in responses to viral infection and disease resistance (Li et al., 2020) (Zhao et al., 2022). These findings raise the possibility that *HvARF12* has undergone adaptive evolution, possibly driven by trade-offs between agronomic performance and stress tolerance.

Taken together, these results reveal diverse evolutionary dynamics within the *ARF* family in barley, shaped by natural selection, domestication, and breeding. The combination of phylogenetic, structural, and *Ka/Ks* analyses provides key insights into functional divergence and adaptation of *ARF* genes in cereal crops.

### TE identification, classification, and their regulatory influence on *ARF* gene expression

Transposable elements (TEs) are mobile genetic elements capable of within the genome, frequently inducing mutations or altering gene expression. Representing a substantial fraction of plant genomes, TEs have been recognized as major contributors to genome evolution, structural variation, and transcriptional regulation (Bennetzen and Wang, 2014; Hirsch and Springer, 2017; Lisch, 2013). To explore the potential *cis*-regulatory effects of TEs on *ARF* gene expression, we surveyed TE insertions within 100 kb upstream and downstream of all *ARF* genes across 76 barley accessions, using a high-confidence TE annotation dataset.

In total, 516 TEs were identified near *ARF* loci, encompassing diverse TE families, with retrotransposons constituting the predominant class (Figure 3A). Interestingly, TE insertions were not randomly distributed: significantly enrichment was observed within the 50 kb upstream of many *HvARF* genes. A secondary enrichment peak occurred around 20 kb upstream of several loci (Figure 3B). Given their proximity to core promoter regions and their established roles in modulating chromatin accessibility and transcription factor binding (Hirsch and Springer, 2017; Lönnig and Saedler, 2002), these upstream TEs are strong candidates for modulating *ARF* gene expression via *cis*-regulatory mechanisms. The observed spatial enrichment pattern suggests non-random TE insertion or retention, possibly reflecting selection for regulatory function. To investigate the potential regulatory impact of nearby TEs, we conducted expression quantitative trait loci (eQTL) mapping (Druka et al., 2010) in a panel of 20 representative genotypes by integrating high-quality pan-genome and pan-transcriptome datasets (Guo et al., 2025). To disentangle the complex sources of expression variation, we partitioned the variance in gene expression into components attributable to genotype, tissue, and their interaction. Among the 26 *ARF* genes, 22 displayed sufficient expression variability and were retained for further analysis.

**Fig. 3.**
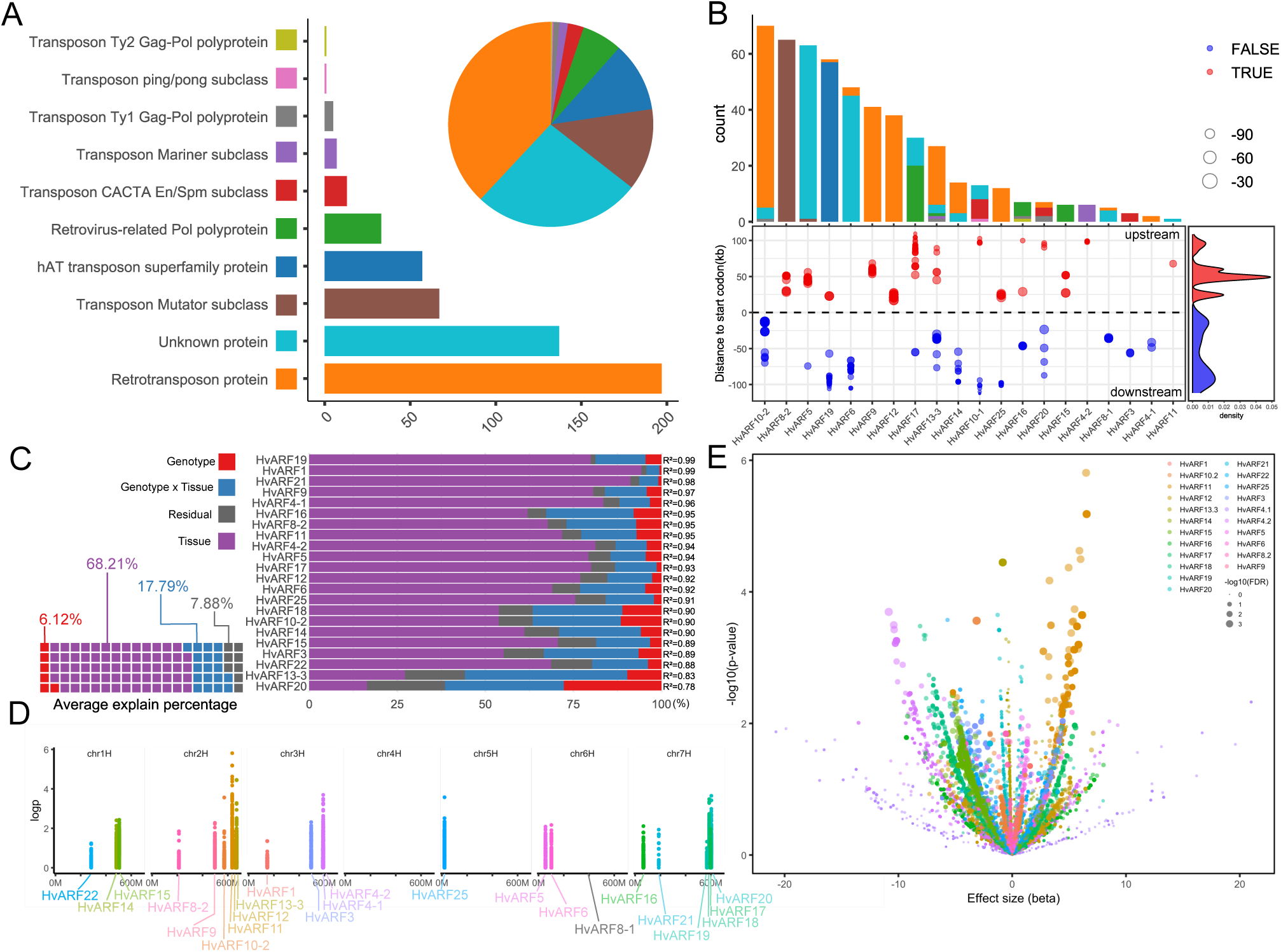
Classification of TEs and eQTL analysis of *ARF* genes. (A) Number and classification of TEs located within 100 kb flanking regions of each *ARF* gene across 76 barley genotypes. (B) TE types, counts, and positional distribution around *ARF* genes. Density plot shows the enrichment of TEs relative to *ARF* gene loci. (C) Proportion of expression variance explained each *ARF* gene by genotype, tissue and genotype × tissue interaction. The waffle chart displays the average contribution of each factor across all genes. (D) *cis*-eQTL signals for *ARF* genes within a ±1 Mb region. (E) Volcano plot of eQTL-associated SNPs. The x-axis represents effect size, and the y-axis shows -log10(P-value).

Variance component analysis revealed that tissue and tissue × genotype interactions were the primary drivers of expression variation, consistent with the known tissue-specific expression profiles of *ARF* genes (Figure 3C). To mitigate tissue effects in subsequent association analyses, we used residual expression values—obtained by regressing out tissue effects—instead of raw expression levels. *Cis*-eQTLs were identified using SNPs located within a 1 Mb window flanking each *ARF* gene. As anticipated, genes with high genotype or genotype-by-tissue explained variance (PVE) —such as *HvARF20*, *HvARF13-3*, *HvARF18*, and *HvARF25*—displayed strong eQTL signals (–logCCP > 3) (Figure 3D, E). Notably, *HvARF25* exhibited a distinct eQTL peak located 23–32 kb upstream of its transcriptional start site, coinciding with the position of a retrotransposon. This suggests that the TE insertion may act as a *cis*-regulatory element modulating *HvARF25* expression in a genotype-dependent manner (Figure 3B, D).

A particularly interesting outlier was *HvARF11*, which showed the most significant eQTL signal despite having low genotype and genotype-by-tissue PVE, and lacking annotated TEs within its vicinity (Figure 3 D, E). This implies the presence of a potent, localized regulatory variant—potentially an unannotated TE insertion, structural variation, or epigenetic modification—contributing substantially to *HvARF11* expression, independent of broader genomic background effects. Overall, TE located near *ARF* genes are likely to play a regulatory role contributing to genotype-specific expression differences.

### Pan-transcriptomic analysis reveals tissue-specific expression and functional networks of *ARF* genes

To elucidate the functional roles and tissue-specific regulatory dynamics of *ARF* genes in barley, we integrated population-scale pan-genomic (Jayakodi et al., 2024) and pan-transcriptomic (Guo et al., 2025) datasets. A total of 22 *ARF* genes with consistently detectable expression across five major tissues in 20 representative genotypes were retained for downstream analyses, minimizing noise from low-abundance transcripts.

*ARF* genes displayed pronounced tissue-specific expression patterns, with the inflorescence exhibiting the highest genotype-dependent expression variability, followed by roots (Figure 4A). Notably, 12 of the 22 *ARF* genes were highly and specifically expressed in developing inflorescences at Waddington stages W6–W7 (Waddington et al., 1983), a critical window for spikelet and floret meristem differentiation. Seven genes showed peak expression in seedling roots, while *HvARF2* and *HvARF6* were predominantly expressed in shoots tissues (Figure 4B). *HvARF22* exhibited unique expression in caryopsis, and no ARF genes demonstrated tissue specificity in embryonic structures such as the embryo, mesocotyl, or seminal root. To further resolve the spatial expression dynamics within inflorescence tissue, we utilized a high-resolution single-cell RNA-seq atlas of barley inflorescence (Morex) (Demesa-Arevalo et al., 2025). This analysis uncovered cell-type-specific expression patterns for individual ARF genes (Figure 4C). *HvARF4-1* and *HvARF4-2* were broadly expressed across diverse cell types, indicating general developmental roles. In contrast, *HvARF14*, *HvARF15*, and *HvARF11* were enriched in vascular-associated tissues, with *HvARF11* being specifically expressed during the transition from triple-spikelet meristem to floret meristem—an essential developmental checkpoint. Additionally, *HvARF6*, *HvARF12*, and *HvARF25* were predominantly expressed in the adaxial epidermis, suggesting possible roles in organ polarity or boundary formation. These temporally and spatially resolved expression profiles support the hypothesis that ARF genes operate in multiple, partially distinct regulatory programs during inflorescence development.

**Fig. 4.**
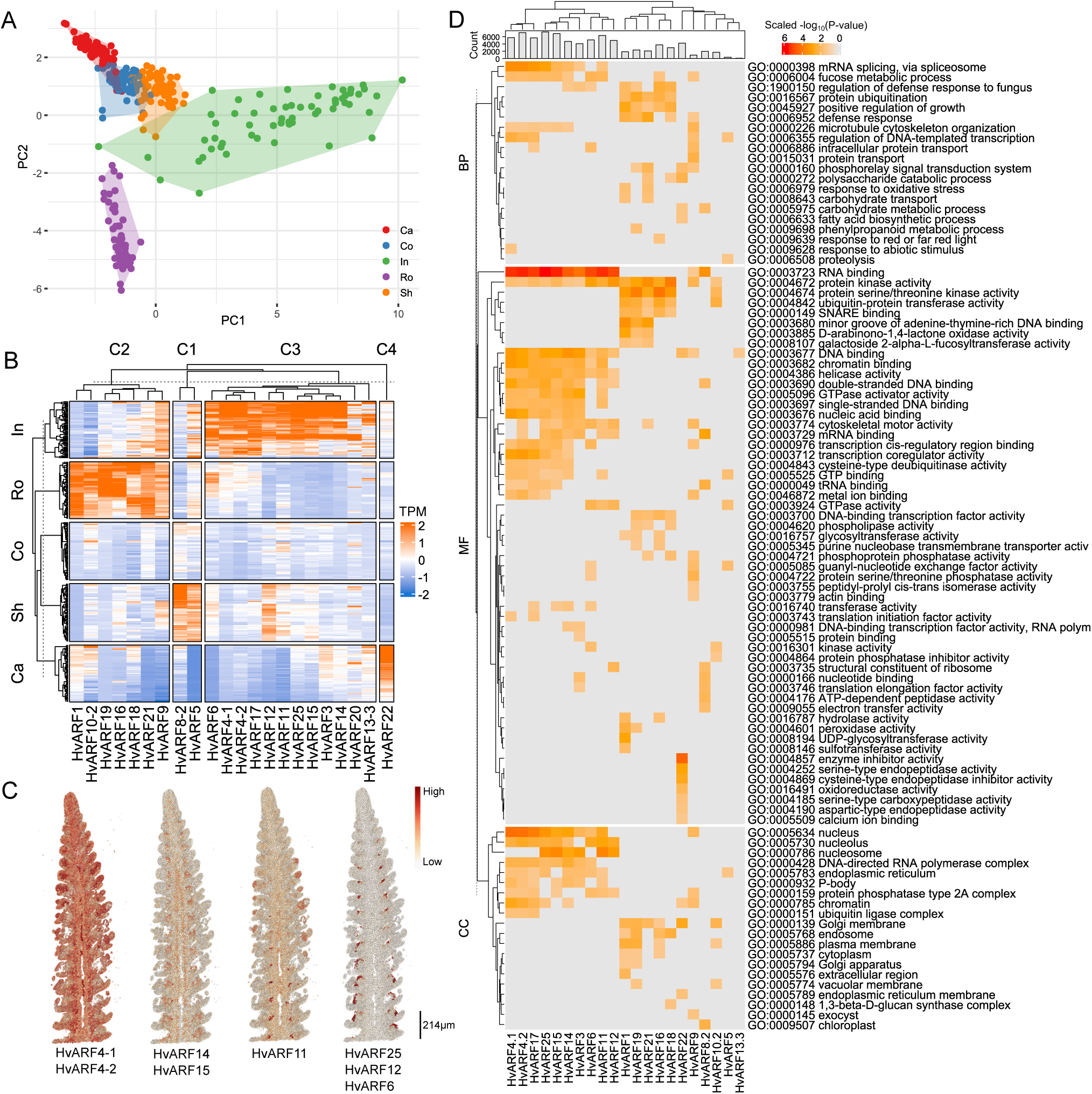
Pan-transcriptome analysis of each *ARF* gene expression. (A) PCA of *ARF* gene expression across all genotype-tissue combinations. PC1 and PC2 explain 41.5% and 24.1% of the total variance, respectively. Colors indicate tissue types; shaded areas highlight genotype-dependent variation within each tissue. (B) Summary heatmap showing expression levels of *ARF* genes across five tissues and 20 genotypes. (C) Single-cell transcriptomic visualization of *ARF* gene expression in barley inflorescence. Color intensity represents relative expression level; scale bar = 214 µm. (D) GO enrichment analysis of gene co-expression modules associated with each *ARF* gene (co-expression threshold ≥ 0.7). Color scale indicates -log_10_(P-value) normalized by Z-score.

To further explore the potential regulatory networks associated with those ARF gene, we conducted a weighted gene co-expression network analysis (WGCNA). Residual expression values (adjusted for genotypic effects) were used as input to focus on tissue-specific transcriptional relationships. For each ARF gene, co-expressed gene modules were identified, and Gene Ontology (GO) enrichment analysis was performed (Figure 4D). As expected for transcription factors, most *HvARF*-associated modules were enriched in functions related to RNA/DNA binding and transcriptional regulation. Notably, *HvARF3*, *HvARF14*, and *HvARF15* were co-expressed with genes involved in fungal defense responses. This observation is consistent with the functional identification of their rice orthologs (*OsARF3*, *OsARF3a*, *OsARF3b*), which have been reported to modulate disease resistance pathways.

Overall, our discovery indicates that *ARF* genes exhibit tissue and cell-type-specific expression patterns and are involved in distinct regulatory modules that govern a range of biological processes—from developmental patterning in inflorescence to biotic stress responses. These results insight a foundational framework for future functional studies and genetic manipulation strategies for optimizing inflorescence architecture or resistance in barley.

### Pangenome-based multi-omics analysis identifies HvARF3 as a key target of selection during barley breeding

To investigate the role of *ARF* gene variation in barley domestication and improvement, we performed selective sweep analyses alongside functional predictions of amino acid (AA) substitutions. Selection signals were examined within genomic regions flanking each *ARF* gene using a population panel comprising wild barley, landraces, and modern cultivars (Figure S3). Among the 26 *ARF* loci, five— *HvARF3*, *HvARF8-2*, *HvARF21*, *HvARF15*, and *HvARF4-1*—exhibited strong selection signatures, with marked allelic differentiation between wild and domesticated accessions (Figure 5A). These findings suggest that these loci may have undergone positive selection due to their contributions to agronomic traits favored during domestication and breeding.

**Fig. 5.**
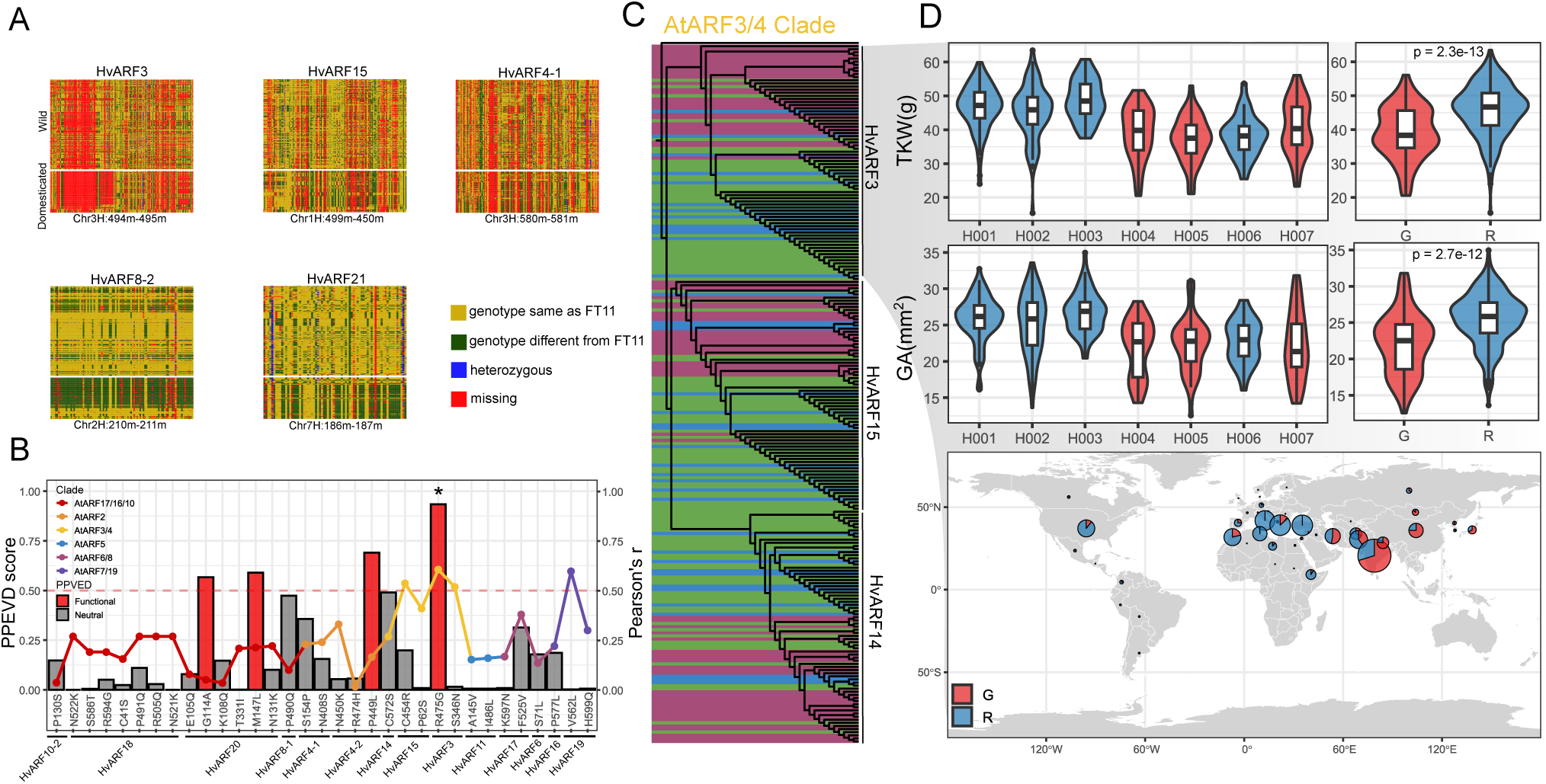
Germplasm analysis of *ARF* genes and haplotype analysis of HvARF3. (A) SNP matrix showing putative selective sweep regions for five *ARF* genes. Different colors indicate nucleotide polymorphisms at each site across genotypes. (B) Bar plot showing PPVED-predicted functional impact scores (left y-axis) and Pearson’s correlation coefficients (right y-axis) for amino acid substitutions across ARF proteins. (C) Phylogenetic trees of HvARF3, HvARF14 and HvARF15, with wild (purple), landrace (blue), and cultivar (green) accessions indicated. (D) Haplotype network and geographic distribution of HvARF3 across the barley diversity panel.

To assess the potential functional relevance of naturally occurring amino acid substitutions, we predicted the effects of all non-synonymous variants (allele frequency > 0.1) using the Plant Protein Variation Effect Detector (PPVED) (Gou et al., 2022). Pearson correlation analyses between amino acid variants and domestication scores identified several candidate residues under selection (Figure 5B). Four variants had PPVED scores > 0.5, indicating a high probability of functional impact, while the remaining substitutions were predicted to be functionally neutral. The most notable protein variant was a non-synonymous substitution in HvARF3 (R475G), which had the highest predicted functional impact across the ARF protein family. HvARF3, along with HvARF14 and HvARF15, belongs to the AtARF3/4 orthologous clade. All three members of this subclade displayed strong selection signals despite being located on different chromosomes, indicating potential convergent selection acting on this regulatory lineage (Figure 5B). Phylogenetic analysis of ARF proteins across the 76 barley accessions further supported this observation. While most ARF clades contained a mixture of wild and domesticated accessions, HvARF3, HvARF14, and HvARF15 formed lineage-specific clusters corresponding to domestication groups (Figure 5C). This convergence of selective sweep signals, phylogenetic divergence, and predicted functional variant highlights the AtARF3/4 subclade—particular HvARF3—as a key target of directional selection linked to agronomic adaptation.

To investigate the phenotypic consequences of *ARF* gene variation, we performed genome-wide association studies (GWAS) focused on key yield components, particularly grain size and thousand-kernel weight (TKW). SNPs within 1 Mb of each *ARF* gene were analyzed using a significance threshold of – logCC(P) > 2. While most associations showed modest effect sizes, consistent with the pleiotropic and conserved roles of *ARF* genes, we observed consistent association peaks near *HvARF3* and *HvARF4-1*, particularly for TKW and grain size (Figure S2 C, Table S4). To validate the association of *HvARF3* with grain traits, we performed haplotype analysis based on SNPs within the gene body and its flanking regulatory regions. Seven major haplotypes were identified and could be classified into two distinct groups based on the R475G substitution: the “R475” allele, predominantly present in European cultivars, and the “G475” allele, more frequently observed in Asian accessions. Accessions harboring the R475 allele exhibited significantly higher grain area (GA) and TKW, two key traits under strong selection during modern barley breeding (Figure 5D). Notably, this positive association between the R475 allele and increased grain size and weight remained significant even after correcting for the *nud* locus (responsible for the naked vs. hulled grain phenotype), indicating that allelic variation at *HvARF3* contributes to grain trait variation independently of hull type (Figure S2B).

To facilitate marker-assisted selection, a derived cleaved amplified polymorphic sequence (dCAPS) marker was developed to discriminate between the R and G alleles at the HvARF3 R475G site (Figure S5). Collectively, this result provides indirect but convincing evidence that *HvARF3* regulates grain traits and has undergone selection during barley improvement.

## Discussion

In this study, we present a novel method of gene family analysis by utilizing abundant, publicly available genomic and transcriptomic resources to comprehensively investigate the *ARF* gene family in barley. Through the integration of multi-omics datasets, we performed a series of systematic analyses including PAV detection, evolutionary dynamics, TE-mediated eQTL mapping, transcriptome analysis, and domestication signal identification. A summary of *ARF* genes in barley has been created by using different data source and methods, which provide insights and clues for future cloning and analysis. Those analyses, which were previously impossible with a single reference genome, demonstrate the advantage of pan-genome, pan-transcriptome, and resequencing data in mining gene family complexity across diverse genotypes in barley.

Recent advances in barley multi-omics resources, including high-resolution pan-genomes and population-wide transcriptomes, offer unprecedented opportunities for gene family research. However, the challenge lies in effective data integration and interpretation. Our study demonstrates that combining these layers enables the discovery of functional alleles and regulatory mechanisms otherwise inaccessible. For instance, the identification of HvARF3^R475G^ as a putative domestication-related variant associated with increased grain size and TKW underscores the value of population-scale pan-genomic approaches. Conventional gene family studies relying on single genomes often fail to capture intraspecies diversity— especially in crops like barley with global distribution and rich germplasm pools. Pan-genomics strategies, in contrast, enable the identification of both structural and nucleotide-level polymorphisms, accelerating trait mapping and gene discovery. Previously, genes such as *Ppd-H1* and *VRN-1* were identified through large-scale germplasm screenings (Santra et al., 2009; Turner et al., 2005). Today, pan-genomic datasets allow for earlier and more targeted discovery. In this study, we revealed geographic stratification of HvARF3 haplotypes—paralleling patterns observed in well-characterized flowering regulators like wheat *Ppd-D1*— highlighting possible regional adaptation and breeding selection.

The *ARF* gene family comprises a moderate number of transcription factors with essential roles in auxin-mediated growth and development. While several *ARFs* have been functionally characterized in *Arabidopsis* and rice, their roles in barley remain underexplored. Phylogenetic analyses revealed lineage-specific expansions and contractions. Notable, the AtARF17/16/10 clade exhibited multiple duplications in barley, particularly within the *HvARF13* subgroup. *HvARF13-3* remains highly conserved, while *HvARF13-1* and *HvARF13-2* show CNVs and sequence divergence—suggesting ongoing sub- functionalization or neofunctionalization (Figure 1B). These variations are more frequent in wild and landrace accessions, implying adaptive relevance in non-domesticated environments. Comparative genomics also revealed notable differences in *ARF* gene retention and loss. For example, rice contains two paralogs, *OsARF23* and *OsARF24* (formerly OsARF1), involved in grain formation via *rice morphology determinant (RMD)* regulation (Li et al., 2014; Waller et al., 2002). These genes have no direct orthologs in barley (Figure 2A), indicating species-specific divergence. Conversely, barley harbors a duplicated *OsARF4* ortholog pair—*HvARF4-1* and *HvARF4-2*—both retaining full-length domain structures. Given *OsARF4’s* known role in negatively regulating grain size in rice (Hu et al., 2018), its barley counterparts represent strong candidates for functional studies.

Despite previous findings that several rice *ARF* genes (e.g., *OsARF3*, *OsARF4*, *OsARF6*, *OsARF14*, *OsARF15*) are associated with grain-related traits (Gu et al., 2023; Hu et al., 2018; Qiao et al., 2021), most *ARF* genes in barley do not show direct expression in the caryopsis (Figure 4B). Instead, their strong expression in inflorescence tissues suggests that these *ARF* genes may regulate grain development indirectly by modulating early reproductive organogenesis and floral architecture, which likely affects final grain size or weight. Another example is *HvARF3*, the selected gene during breeding, it shows high gene expression levels in the developing inflorescence while no expression in grain. In the single-cell transcriptome dataset of barley inflorescences, *HvARF3* is specifically expressed in dividing cells, suggesting a potential role in regulating cell division and meristem maintenance during early stages of inflorescence development. But according to GWAS haplotype analysis and performance of its ortholog *OsARF3*, it most likely affects grain size and grain traits, too. Interestingly, the favorable haplotypes of *HvARF3* are rare in Asian barley accessions. This may be attributable to a regional genetic bottleneck or local adaptation, which limited the introgression of the advantageous haplotype. Alternatively, certain *HvARF3* alleles may exhibit functional trade-offs similar to its rice ortholog *OsARF3*, which has been shown to mediate a balance between heat tolerance and pathogen resistance (Gu et al., 2023). Furthermore, the *Ka*/*Ks* rate showed that *HvARF3* was higher in Asian accessions, indicating that the purifying selection was not as strong as that in Europe. Thus, environmental difference between European and Asian growth conditions could have shaped regional selection pressures acting on different *HvARF3* variants and reflect different breeding strategy.

In conclusion, our integrative analysis highlights the critical role of *ARF* genes in barley development and demonstrates the utility of multi-omics strategies in gene family research. By resolving patterns of structural variation, expression regulation, and selection, we identify promising candidates like *HvARF3* for targeted breeding. These findings contribute not only to our understanding of auxin signaling but also to practical efforts aimed at improving grain yield and adaptability in barley. The dCAPS marker developed for the HvARF3^R475^ variant will facilitate marker-assisted selection in future breeding programs.

## Methods

### Identification of ARF genes in 76 barley pan-genome accessions

Genomic and annotated protein/coding sequences for 76 barley pan-genome accessions were retrieved from a previously published study (Jayakodi et al., 2024). To comprehensive identify members of the ARF protein family, all candidate proteins containing ARF-related domains—including PTHR31384 from PANTHER; PF02362, PF06507, and PF02309 from Pfam, along with the B3 (PS50863) and PB1 (PS51745) domains—were identified using *HMMER* v3.1b2 (Finn et al., 2011) with an E-value threshold of < 1e−5. Domain annotations were verified by comparing against known ARF proteins from rice and *Arabidopsis* using *BLAST+* v2.13.0 (Camacho et al., 2009). Identified ARF proteins were named based on their phylogenetic relationships to rice ARFs, using the barley cultivar Barke as the reference genome. To determine presence/absence variations (PAVs) and copy number variations (CNVs), we extracted 5 kb flanking sequences of loci where annotated ARF genes were missing using *Bedtools* v2.29.2 (Quinlan and Hall, 2010). These regions were aligned to the Morex reference genome using *Minimap2* v2.16 (Li, 2018). Large deletions were visualized using *MUMmer* v3.23 (Delcher et al., 2003), while smaller deletions were confirmed through BLAST alignments.

### Phylogenetic tree construction and *Ka*/*Ks* calculation

A total of 1,911 ARF protein sequences from the pan-genome were aligned using MAFFT v7.490 (Katoh and Standley, 2013). Phylogenetic relationships were inferred using *IQ-TREE* v2.2.2.6 (Nguyen et al., 2015) with the maximum likelihood method under default parameters. For interspecies comparison, ARF protein sequences from barley (cv. Barke), rice (cv. Nipponbare), and *Arabidopsis* (Col-0) were used to construct a combined phylogenetic tree using the same procedure. Conserved motifs were identified using MEME v5.3.3 (Bailey et al., 2015). Phylogenetic trees and conserved domain architectures were visualized and edited using the iTOL platform (https://itol.embl.de/). To assess selection pressures, pairwise nonsynonymous to synonymous substitution rate ratios (*Ka/Ks*) were calculated using *ParaAT2.0*, based on aligned ARF ortholog sequences from the 76 barley genotypes. Statistical significance of *Ka/Ks* differences between groups (e.g., wild vs. domesticated accessions) was tested using a two-tailed Student’s t-test.

### Transcriptome data processing and co-expression analysis

Raw transcriptome data from five tissues—caryopsis, coleoptile, inflorescence, root, and shoot—across 20 barley cultivars were retrieved from the recently published barley pan-transcriptome dataset (accession: PRJEB64639). Transcript abundance was quantified using Kallisto v0.46.1 (Bray et al., 2016), yielding TPM values and raw read counts. To minimize genotype-specific effects in the functional analysis of *ARF* genes, genotype-corrected residual expression values (*Residual_G_*) were calculated (see below) and used for downstream analyses. Weighted gene co-expression network analysis (WGCNA) was performed using the *WGCNA* R package with default parameters and a correlation threshold of 0.7. Gene Ontology (GO) enrichment was conducted using *clusterprofiler* v3.2.1 (Yu et al., 2012), with GO annotations sourced from the IPK barley database (https://galaxy-web.ipk-gatersleben.de/). Single-cell expression patterns of *ARF* genes were extracted from a recently published barley inflorescence single-cell transcriptomic resource BARVISTA (https://www.plabipd.de/projects/hannah_demo/BARVISTA/tissue_no_download.html) (Demesa-Arevalo et al., 2025).

### Proportion of variance explained (PVE) and residual calculation

To quantify the contribution of genotype and tissue to gene expression variation, a linear mixed model (LMM) was fitted for each ARF gene using the following equation:

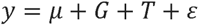

Where *y* represents the expression value (TPM), 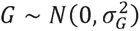) is the random genotype effect, 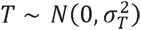 is the tissue effect, and E is the residual error. The proportion of variance explained by genotype and tissue was calculated as:

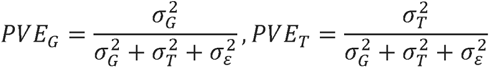

Genotype- and tissue-corrected residuals were then extracted using:

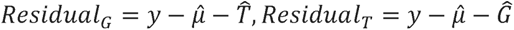

All calculations were performed in R using *lme4* package (Bates et al., 2015).

### TE identification and *cis*-eQTL analysis

TEs were identified in 76 barley genomes using *panHiTE* (Hu et al., 2025), and further confirmed using TE annotations from the IPK panBarlex database (https://panbarlex.ipk-gatersleben.de/#). Only high confidence TEs present in both of datasets were retained. To determine TE insertions near ARF genes, 100 kb flanking regions were analyzed using *bedtools*. For cis-eQTL analysis, 1 Mb upstream and downstream of each ARF gene was defined as the cis-regulatory region. *Residual_T_* values were used as gene expression input in *MatrixEQTL* v2.3 (Shabalin, 2012) under default settings.

### Selective sweep detection

To explore potential domestication-related selection signals, raw genomic data were obtained from a recent barley population study (Guo et al., 2024). A 1 Mb window surrounding each *ARF* gene was analyzed. Genomic regions were extracted using *bcftools* v1.15.1, and downstream processing and visualization were carried out using Perl under the guidance of the original study’s first author (Guo et al., 2024).

### Variant calling and amino acid substitution prediction

Whole-genome assemblies of the 76 barley accessions were re-mapped to the Morex V2 reference genome (Figure, S4). Genomes were split into 1 kb fragments using *seqkit* v0.9.1 (Shen et al., 2016) to simulate FASTQ reads. Variant calling was performed using *BWA* v0.7.17 (Li, 2013), *SAMtools* v1.16.1(Li et al., 2009), and *BCFtools* v1.15.1. SNPs and InDels were annotated using *SnpEff* (Cingolani et al., 2012). Amino acid substitutions in ARF proteins were predicted with *PPVED* v1.0 (Gou et al., 2022). Correlation analysis between substitution scores and domestication degrees were calculated using Pearson’s correlation coefficient in R.

### Genome-wide association study (GWAS) and haplotype analysis

GWAS was performed to identify genomic regions associated with 16 agronomic traits (awn length; culm dry weight; fertility rate; final spikelet number; grain area; grain length; grain number per spikelet; grain weight; grain width; heading date; plant height; potential spikelet number; pre-anthesis tip degeneration; spike length; spike weight; thousand kernels weight). All phenotypic data are collected at IPK Gatersleben in 2018, 2019 and 2020. Total 442 representative genotypes were used in this study; population structure was fixed. Some parts of the phenotypic data have been published in previous work (Huang et al., 2023; Kamal et al., 2022). After quality filtering, 33,138,758 SNPs (minor allele frequency > 0.1; missing rate < 10%) were retained. GWAS was conducted using *PLINK* v1.90b6.9 (Purcell et al., 2007) and *GEMMA* v0.98.5 (Team et al., 2024). Haplotype blocks around *ARF* genes (±1 Mb) were analyzed using pipeline shared in github (https://github.com/TKNsama/R). Since some phenotypic data are unpublished, only trait-associated SNPs proximal to *ARF* loci are reported here.

### dCAPS marker design and validation

To genotype the functional HvARF3 R475G variant, a dCAPS marker was developed targeting a SNP located at chr3H:494973445 (based on the MorexV2 reference genome). Primers (Supplementary data 5) were designed using the online tools dCAPS Finder 2.0 and Primer3, incorporating a mismatch to generate a specific restriction site for the enzyme *DdeI* (New England Biolabs Inc). PCR amplification was carried out using *Taq* DNA Polymerase (Qiagen, Cat. No. 201203) following the manufacturer’s recommended protocol. Amplified products were subsequently digested with *DdeI* and separated on a 2.5% agarose gel.

## Data Availability

76 pan-genome sequence and annotation data are from previous publication (Jayakodi et al., 2024); Genotypic data of wild barley/ domesticated barley are from previous publication (Guo et al., 2024); Genotypic data of GWAS will be released in upcoming publication; some phenotypic data (FSN, GY, HD, MYP, PH, PSN, SA) are from previous publication (Kamal et al., 2022), while rest of them will be released in upcoming publication. The re-mapping VCF files created by this study has been deposited at Zenodo (https://zenodo.org/records/16778621).

## Author contributions

KT: funding acquisition, conceptualization, data analysis, visualization, and writing — original draft preparation. ZG: conceptualization, investigation, validation, and writing — review and editing. TS: funding acquisition, supervision, and writing — review and editing.

## Acknowledgments

We thank the public databases for providing valuable resources that supported this study. We also thank Roop Kamal, Yongyu Huang for phenotypic data collection. This work was supported by the Chinese Scholarship Council (CSC) to K.T., and by the Leibniz Institute of Plant Genetics and Crop Plant Research (IPK) through infrastructure and core funding.

## Reference

Aravind, L., Watanabe, H., Lipman, D.J., Koonin, E.V., 2000. Lineage-specific loss and divergence of functionally linked genes in eukaryotes. Proceedings of the National Academy of Sciences 97(21), 11319–11324.

Bailey, T.L., Johnson, J., Grant, C.E., Noble, W.S., 2015. The MEME suite. Nucleic acids research 43(W1), W39–W49.

Bates, D., Mächler, M., Bolker, B., Walker, S., 2015. Fitting linear mixed-effects models using lme4. Journal of statistical software 67, 1–48.

Bennetzen, J.L., Wang, H., 2014. The contributions of transposable elements to the structure, function, and evolution of plant genomes. Annual review of plant biology 65(1), 505–530.

Boden, S., McIntosh, R., Uauy, C., Krattinger, S.G., Dubcovsky, J., Rogers, W.J., Xia, X., Badaeva, E., Bentley, A., Brown-Guedira, G., 2023. Updated guidelines for gene nomenclature in wheat. Theoretical and Applied Genetics 136(4), 72.

Bray, N.L., Pimentel, H., Melsted, P., Pachter, L., 2016. Near-optimal probabilistic RNA-seq quantification. Nature biotechnology 34(5), 525–527.

Camacho, C., Coulouris, G., Avagyan, V., Ma, N., Papadopoulos, J., Bealer, K., Madden, T.L., 2009. BLAST+: architecture and applications. BMC bioinformatics 10, 1–9.

Cancé, C., Martin-Arevalillo, R., Boubekeur, K., Dumas, R., 2022. Auxin response factors are keys to the many auxin doors. New Phytologist 235(2), 402–419.

Chandler, J.W., 2016. Auxin response factors. Plant, cell & environment 39(5), 1014–1028.

Choi, H.-S., Seo, M., Cho, H.-T., 2018. Two TPL-binding motifs of ARF2 are involved in repression of auxin responses. Frontiers in Plant Science 9, 372.

Cingolani, P., Platts, A., Wang, L.L., Coon, M., Nguyen, T., Wang, L., Land, S.J., Lu, X., Ruden, D.M., 2012. A program for annotating and predicting the effects of single nucleotide polymorphisms, SnpEff: SNPs in the genome of Drosophila melanogaster strain w1118; iso-2; iso-3. fly 6(2), 80–92.

Coulter, M., Entizne, J.C., Guo, W., Bayer, M., Wonneberger, R., Milne, L., Schreiber, M., Haaning, A., Muehlbauer, G.J., McCallum, N., 2022. BaRTv2: a highly resolved barley reference transcriptome for accurate transcript-specific RNA-seq quantification. The Plant Journal 111(4), 1183–1202.

Delcher, A.L., Salzberg, S.L., Phillippy, A.M., 2003. Using MUMmer to identify similar regions in large sequence sets. Current protocols in bioinformatics(1), 10.13.11–10.13.18.

Demesa-Arevalo, E., Dörpholz, H., Vardanega, I., Maika, J.E., Pineda-Valentino, I., Eggels, S., Lautwein, T., Köhrer, K., Schnurbusch, T., von Korff, M., Usadel, B., Simon, R., 2025. Imputation integrates single-cell and spatial gene expression data to resolve transcriptional networks in barley shoot meristem development. bioRxiv, 2025.2005.2009.653223. 10.1101/2025.05.09.653223.

Druka, A., Potokina, E., Luo, Z., Jiang, N., Chen, X., Kearsey, M., Waugh, R., 2010. Expression quantitative trait loci analysis in plants. Plant biotechnology journal 8(1), 10–27.

Finet, C., Berne-Dedieu, A., Scutt, C.P., Marlétaz, F., 2013. Evolution of the ARF gene family in land plants: old domains, new tricks. Molecular Biology and Evolution 30(1), 45–56.

Finn, R.D., Clements, J., Eddy, S.R., 2011. HMMER web server: interactive sequence similarity searching. Nucleic acids research 39(suppl_2), W29–W37.

Gao, J., Zhang, L., Du, H., Dong, Y., Zhen, S., Wang, C., Wang, Q., Yang, J., Zhang, P., Zheng, X., 2023. An ARF24-ZmArf2 module influences kernel size in different maize haplotypes. Journal of Integrative Plant Biology 65(7), 1767–1781.

Gidhi, A., Kumar, M., Mukhopadhyay, K., 2021. The auxin response factor gene family in wheat (Triticum aestivum L.): Genome-wide identification, characterization and expression analyses in response to leaf rust. South African Journal of Botany 140, 312–325.

Gou, X., Feng, X., Shi, H., Guo, T., Xie, R., Liu, Y., Wang, Q., Li, H., Yang, B., Chen, L., 2022. PPVED: A machine learning tool for predicting the effect of single amino acid substitution on protein function in plants. Plant Biotechnology Journal 20(7), 1417–1431.

Gu, X., Si, F., Feng, Z., Li, S., Liang, D., Yang, P., Yang, C., Yan, B., Tang, J., Yang, Y., 2023. The OsSGS3-tasiRNA-OsARF3 module orchestrates abiotic-biotic stress response trade-off in rice. Nature Communications 14(1), 4441.

Guo, N., Wang, Y., Chen, W., Tang, S., An, R., Wei, X., Hu, S., Tang, S., Shao, G., Jiao, G., 2022. Fine mapping and target gene identification of qSE4, a QTL for stigma exsertion rate in rice (Oryza sativa L.). Frontiers in Plant Science 13, 959859.

Guo, W., Schreiber, M., Marosi, V.B., Bagnaresi, P., Jørgensen, M.E., Braune, K.B., Chalmers, K., Chapman, B., Dang, V., Dockter, C., 2025. A barley pan-transcriptome reveals layers of genotype-dependent transcriptional complexity. Nature Genetics, 1–10.

Guo, Y., Jayakodi, M., Himmelbach, A., Ben-Yosef, E., Davidovich, U., David, M., Hartmann-Shenkman, A., Kislev, M., Fahima, T., Schuenemann, V.J., Reiter, E., Krause, J., Steffenson, B.J., Stein, N., Weiss, E., Mascher, M., 2024. A haplotype-based evolutionary history of barley domestication. bioRxiv, 2024.2012.2018.628695. 10.1101/2024.12.18.628695.

Hirsch, C.D., Springer, N.M., 2017. Transposable element influences on gene expression in plants. Biochimica et Biophysica Acta (BBA)-Gene Regulatory Mechanisms 1860(1), 157–165.

Hu, K., Xu, M., Wang, J., 2025. panHiTE: a comprehensive and accurate pipeline for TE detection in large-scale population genomes. bioRxiv, 2025.2002. 2015.638472.

Hu, Z., Lu, S.J., Wang, M.J., He, H., Sun, L., Wang, H., Liu, X.H., Jiang, L., Sun, J.L., Xin, X., 2018. A novel QTL qTGW3 encodes the GSK3/SHAGGY-like kinase OsGSK5/OsSK41 that interacts with OsARF4 to negatively regulate grain size and weight in rice. Molecular Plant 11(5), 736–749.

Huang, J., Li, Z., Zhao, D., 2016. Deregulation of the Os miR160 target gene OsARF18 causes growth and developmental defects with an alteration of auxin signaling in rice. Scientific Reports 6(1), 29938.

Huang, Y., Kamal, R., Shanmugaraj, N., Rutten, T., Thirulogachandar, V., Zhao, S., Hoffie, I., Hensel, G., Rajaraman, J., Moya, Y.A.T., Hajirezaei, M.R., Himmelbach, A., Poursarebani, N., Lundqvist, U., Kumlehn, J., Stein, N., von Wiren, N., Mascher, M., Melzer, M., Schnurbusch, T., 2023. A molecular framework for grain number determination in barley. Sci Adv 9(9), eadd0324.

Jayakodi, M., Lu, Q., Pidon, H., Rabanus-Wallace, M.T., Bayer, M., Lux, T., Guo, Y., Jaegle, B., Badea, A., Bekele, W., 2024. Structural variation in the pangenome of wild and domesticated barley. Nature, 1–9.

Jayakodi, M., Padmarasu, S., Haberer, G., Bonthala, V.S., Gundlach, H., Monat, C., Lux, T., Kamal, N., Lang, D., Himmelbach, A., 2020. The barley pan-genome reveals the hidden legacy of mutation breeding. Nature 588(7837), 284–289.

Jia, M., Li, Y., Wang, Z., Tao, S., Sun, G., Kong, X., Wang, K., Ye, X., Liu, S., Geng, S., 2021. TaIAA21 represses TaARF25-mediated expression of TaERFs required for grain size and weight development in wheat. The Plant Journal 108(6), 1754–1767.

Kamal, R., Muqaddasi, Q.H., Zhao, Y., Schnurbusch, T., 2022. Spikelet abortion in six-rowed barley is mainly influenced by final spikelet number, with potential spikelet number acting as a suppressor trait. J Exp Bot 73(7), 2005–2020.

Katoh, K., Standley, D.M., 2013. MAFFT multiple sequence alignment software version 7: improvements in performance and usability. Molecular biology and evolution 30(4), 772–780.

Kimura, M., 1985. The neutral theory of molecular evolution. Cambridge university press.

Li, G., Liang, W., Zhang, X., Ren, H., Hu, J., Bennett, M.J., Zhang, D., 2014. Rice actin-binding protein RMD is a key link in the auxin–actin regulatory loop that controls cell growth. Proceedings of the National Academy of Sciences 111(28), 10377–10382.

Li, H., 2013. Aligning sequence reads, clone sequences and assembly contigs with BWA-MEM. arXiv preprint arXiv:1303.3997.

Li, H., 2018. Minimap2: pairwise alignment for nucleotide sequences. Bioinformatics 34(18), 3094–3100.

Li, H., Handsaker, B., Wysoker, A., Fennell, T., Ruan, J., Homer, N., Marth, G., Abecasis, G., Durbin, R., Subgroup, G.P.D.P., 2009. The sequence alignment/map format and SAMtools. bioinformatics 25(16), 2078–2079.

Li, Y., Han, S., Qi, Y., 2023. Advances in structure and function of auxin response factor in plants. Journal of Integrative Plant Biology 65(3), 617–632.

Li, Y., Li, J., Chen, Z., Wei, Y., Qi, Y., Wu, C., 2020. OsmiR167a-targeted auxin response factors modulate tiller angle via fine-tuning auxin distribution in rice. Plant Biotechnology Journal 18(10), 2015–2026.

Lisch, D., 2013. How important are transposons for plant evolution? Nature reviews genetics 14(1), 49–61.

Lister, D.L., Jones, H., Oliveira, H.R., Petrie, C.A., Liu, X., Cockram, J., Kneale, C.J., Kovaleva, O., Jones, M.K., 2018. Barley heads east: Genetic analyses reveal routes of spread through diverse Eurasian landscapes. PloS one 13(7), e0196652.

Liu, P.P., Montgomery, T.A., Fahlgren, N., Kasschau, K.D., Nonogaki, H., Carrington, J.C., 2007. Repression of AUXIN RESPONSE FACTOR10 by microRNA160 is critical for seed germination and post-germination stages. The Plant Journal 52(1), 133–146.

Liu, X., Zhang, H., Zhao, Y., Feng, Z., Li, Q., Yang, H.Q., Luan, S., Li, J., He, Z.H., 2013. Auxin controls seed dormancy through stimulation of abscisic acid signaling by inducing ARF-mediated ABI3 activation in Arabidopsis. Proceedings of the National Academy of Sciences 110(38), 15485–15490.

Lönnig, W.-E., Saedler, H., 2002. Chromosome rearrangements and transposable elements. Annual review of genetics 36(1), 389–410.

Mallory, A.C., Bartel, D.P., Bartel, B., 2005. MicroRNA-directed regulation of Arabidopsis AUXIN RESPONSE FACTOR17 is essential for proper development and modulates expression of early auxin response genes. The Plant Cell 17(5), 1360–1375.

Man, Q.C., Wang, Y.C., Gao, S.J., Gao, Z.C., Peng, Z.P., Cui, J.H., 2025. Pan-genome analysis and expression verification of the maize ARF gene family. Frontiers in Plant Science 15, 1506853.

Marks, R.A., Hotaling, S., Frandsen, P.B., VanBuren, R., 2021. Representation and participation across 20 years of plant genome sequencing. Nature plants 7(12), 1571–1578.

Marroni, F., Pinosio, S., Morgante, M., 2014. Structural variation and genome complexity: is dispensable really dispensable? Current Opinion in Plant Biology 18, 31–36.

McSteen, P., 2010. Auxin and monocot development. Cold Spring Harbor perspectives in biology 2(3), a001479.

Nguyen, L.-T., Schmidt, H.A., Von Haeseler, A., Minh, B.Q., 2015. IQ-TREE: a fast and effective stochastic algorithm for estimating maximum-likelihood phylogenies. Molecular biology and evolution 32(1), 268–274.

Okushima, Y., Overvoorde, P.J., Arima, K., Alonso, J.M., Chan, A., Chang, C., Ecker, J.R., Hughes, B., Lui, A., Nguyen, D., 2005. Functional genomic analysis of the AUXIN RESPONSE FACTOR gene family members in Arabidopsis thaliana: unique and overlapping functions of ARF7 and ARF19. The Plant Cell 17(2), 444–463.

Prigge, M.J., Morffy, N., de Neve, A., Szutu, W., Abraham-Juárez, M.J., McAllister, T., Jones, H., Johnson, K., Do, N., Lavy, M., 2025. Comparative mutant analyses reveal a novel mechanism of ARF regulation in land plants. Nature Plants, 1-15.

Purcell, S., Neale, B., Todd-Brown, K., Thomas, L., Ferreira, M.A., Bender, D., Maller, J., Sklar, P., De Bakker, P.I., Daly, M.J., 2007. PLINK: a tool set for whole-genome association and population-based linkage analyses. The American journal of human genetics 81(3), 559–575.

Qiao, J., Jiang, H., Lin, Y., Shang, L., Wang, M., Li, D., Fu, X., Geisler, M., Qi, Y., Gao, Z., 2021. A novel miR167a-OsARF6-OsAUX3 module regulates grain length and weight in rice. Molecular Plant 14(10), 1683–1698.

Quinlan, A.R., Hall, I.M., 2010. BEDTools: a flexible suite of utilities for comparing genomic features. Bioinformatics 26(6), 841–842.

Santra, D.K., Santra, M., Allan, R., Campbell, K., Kidwell, K., 2009. Genetic and molecular characterization of vernalization genes Vrn-A1, Vrn-B1, and Vrn-D1 in spring wheat germplasm from the Pacific Northwest region of the USA. Plant breeding 128(6), 576–584.

Shabalin, A.A., 2012. Matrix eQTL: ultra fast eQTL analysis via large matrix operations. Bioinformatics 28(10), 1353–1358.

Shen, C., Wang, S., Bai, Y., Wu, Y., Zhang, S., Chen, M., Guilfoyle, T.J., Wu, P., Qi, Y., 2010. Functional analysis of the structural domain of ARF proteins in rice (Oryza sativa L.). Journal of experimental botany 61(14), 3971–3981.

Shen, W., Le, S., Li, Y., Hu, F., 2016. SeqKit: a cross-platform and ultrafast toolkit for FASTA/Q file manipulation. PloS one 11(10), e0163962.

Team, G., Mesnard, T., Hardin, C., Dadashi, R., Bhupatiraju, S., Pathak, S., Sifre, L., Rivière, M., Kale, M.S., Love, J., 2024. Gemma: Open models based on gemini research and technology. arXiv preprint arXiv:2403.08295.

Tombuloglu, H., 2019. Genome-wide analysis of the auxin response factors (ARF) gene family in barley (Hordeum vulgare L.). Journal of Plant Biochemistry and Biotechnology 28(1), 14–24.

Turner, A., Beales, J., Faure, S., Dunford, R.P., Laurie, D.A., 2005. The pseudo-response regulator Ppd-H1 provides adaptation to photoperiod in barley. Science 310(5750), 1031–1034.

Waddington, S., Cartwright, P., Wall, P., 1983. A quantitative scale of spike initial and pistil development in barley and wheat. Annals of Botany 51(1), 119–130.

Waller, F., Furuya, M., Nick, P., 2002. OsARF1, an auxin response factor from rice, is auxin-regulated and classifies as a primary auxin responsive gene. Plant Molecular Biology 50, 415–425.

Wang, D., Pei, K., Fu, Y., Sun, Z., Li, S., Liu, H., Tang, K., Han, B., Tao, Y., 2007. Genome-wide analysis of the auxin response factors (ARF) gene family in rice (Oryza sativa). Gene 394(1-2), 13–24.

Wang, H., Huang, Y., Li, Y., Cui, Y., Xiang, X., Zhu, Y., Wang, Q., Wang, X., Ma, G., Xiao, Q., 2024. An ARF gene mutation creates flint kernel architecture in dent maize. Nature Communications 15(1), 2565.

Xing, H., Pudake, R.N., Guo, G., Xing, G., Hu, Z., Zhang, Y., Sun, Q., Ni, Z., 2011. Genome-wide identification and expression profiling of auxin response factor (ARF) gene family in maize. BMC genomics 12, 1–13.

Yu, G., Wang, L.G., Han, Y., He, Q.Y., 2012. clusterProfiler: an R package for comparing biological themes among gene clusters. Omics: a journal of integrative biology 16(5), 284–287.

Zhao, Z.X., Yin, X.X., Li, S., Peng, Y.T., Yan, X.L., Chen, C., Hassan, B., Zhou, S.X., Pu, M., Zhao, J.H., 2022. miR167d-ARF s Module Regulates Flower Opening and Stigma Size in Rice. Rice 15(1), 40.

