## Supplemental Figure for "Pan-genome Analysis Reveals Hidden Diversity and Selection Signatures of Auxin Response Factors (ARFs) Associated with Breeding in Barley"

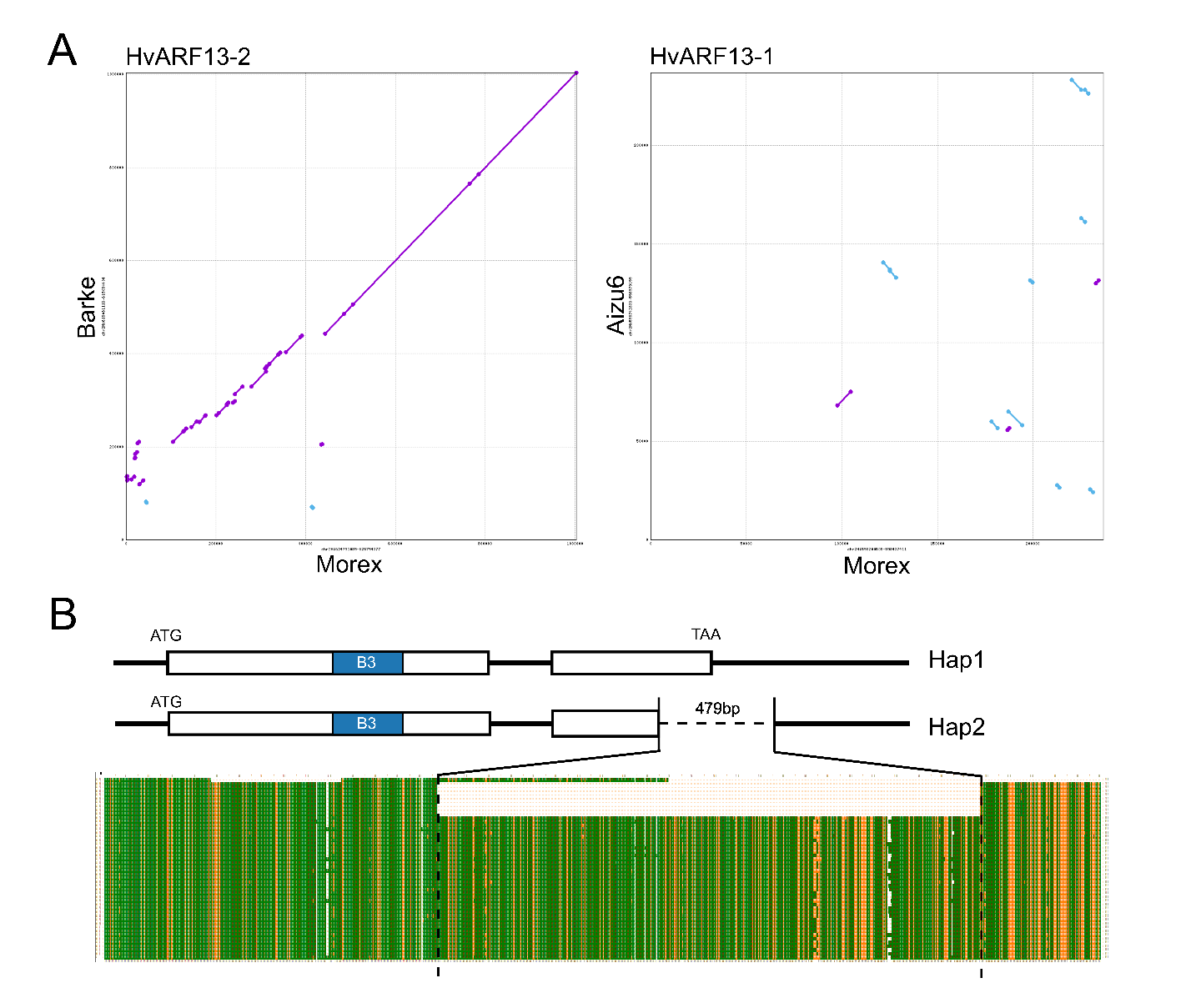

**Supplementary figure 1:** **Genomic variation surrounding *HvARF13-1* and *HvARF13-2*.** (A) Genomic sequence alignment of regions flanking *HvARF13-1* and *HvARF13-2* compared to the Morex reference genome, visualized using MUMmer. (B) Gene structure and BLAST alignment of *HvARF13-2*. The top sequence shows the coding region from cultivar Barke. "Hap1" represents a complete gene structure (e.g., Barke), while "Hap2" corresponds to an incomplete structure with a 479-bp deletion (e.g., Morex). Different colors indicate distinct alignment matches.

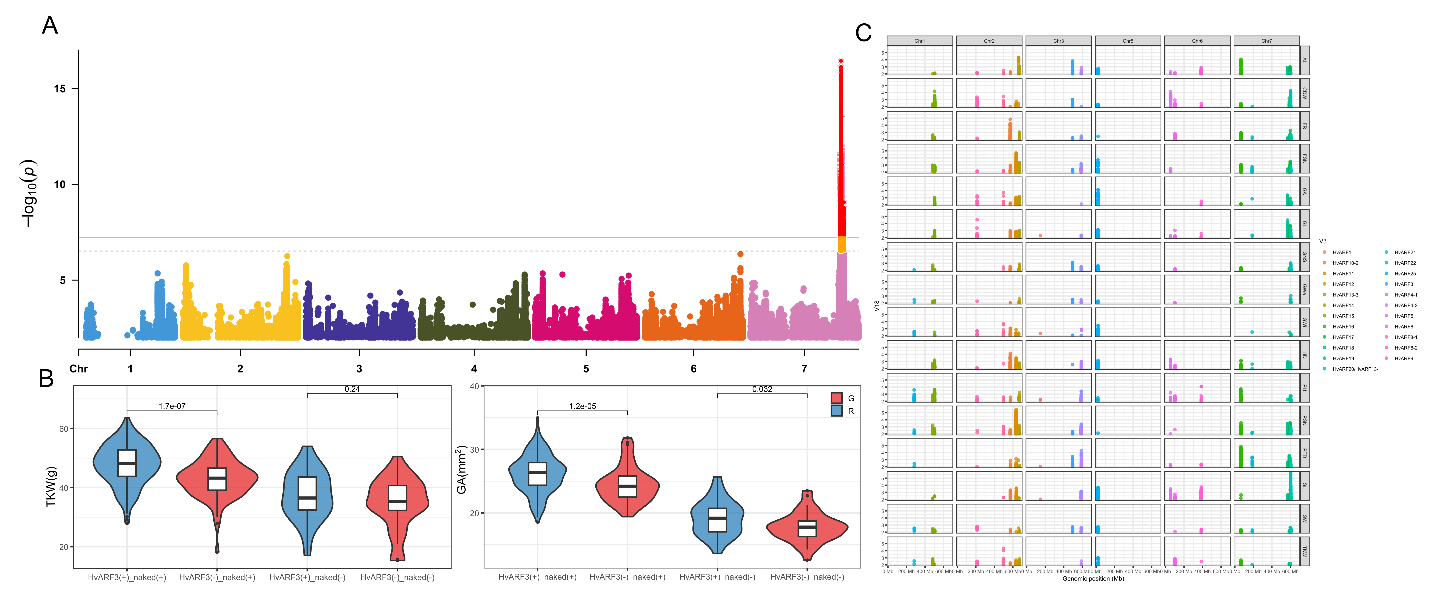

**Supplementary figure 2:** **GWAS panel and additional haplotype analysis.** (A) Manhattan plot of GWAS results for TKW; the most significant association corresponds to the *nud* locus. (B) Two-locus haplotype analysis of the major-effect gene *nud* and *HvARF3*. Statistical significance was evaluated using Student’s *t*-test. (C) GWAS results for multiple traits. Only association peaks located near *ARF* genes are shown. AL, awn length; CDW, culm dry weight; FR, fertility rate; FSN, final spike number; GA, grain area; GL, grain length; GNS, grain number per spike; GWe, grain weight; HD, heading date; PH, plant height; PSN, potential spikelet number; PTD, pre-anthesis degeneration; SL, spike length; SW, spike weight; TKW, thousand kernels weight.

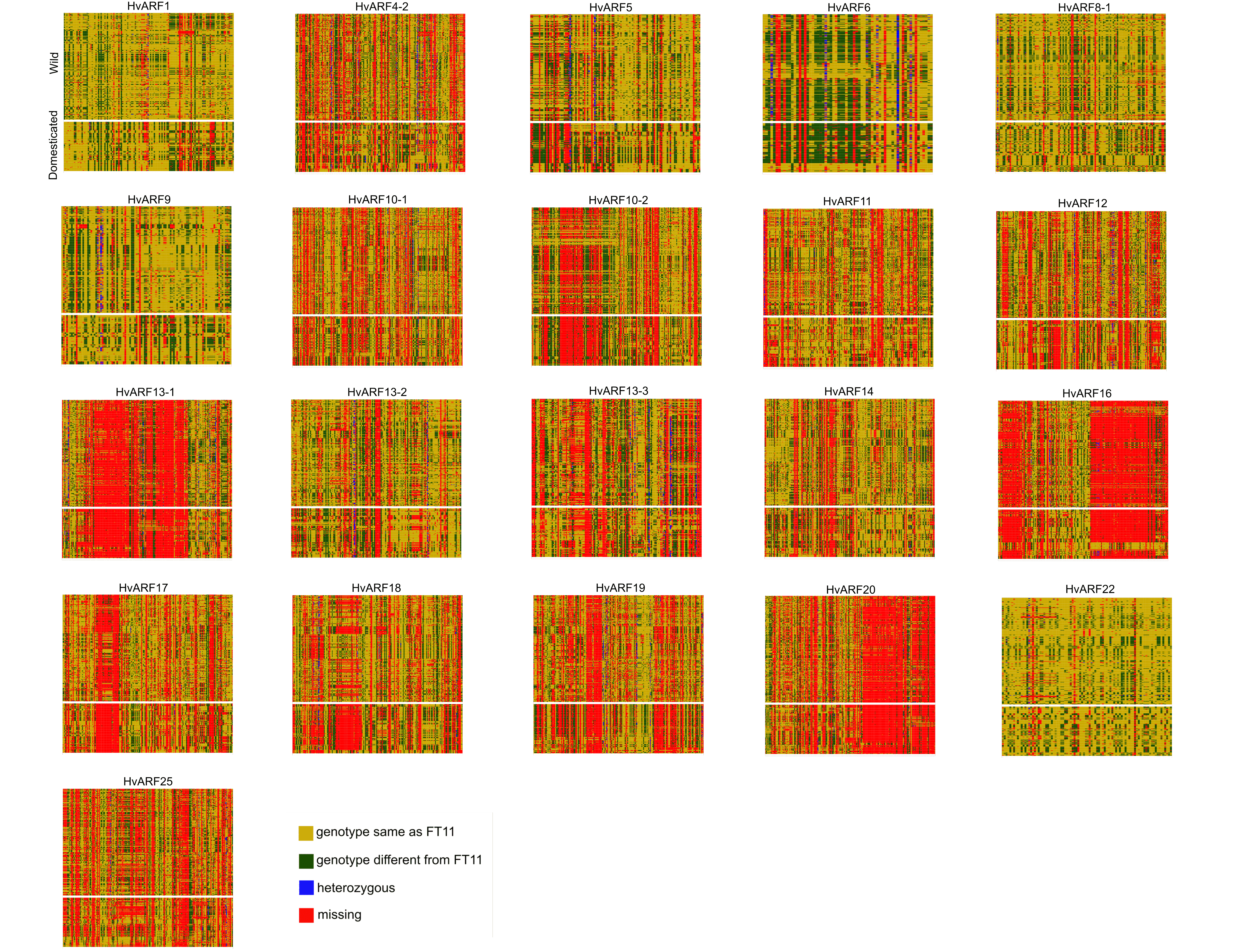

**Supplementary figure 3: Selective sweep regions surrounding *ARF* genes in barley.** Each region displays a 1 Mb genomic region flanking an *ARF* gene, based on the FT11 reference genome. Colors indicate different levels of nucleotide polymorphism, highlighting potential selective sweep signals across the population.

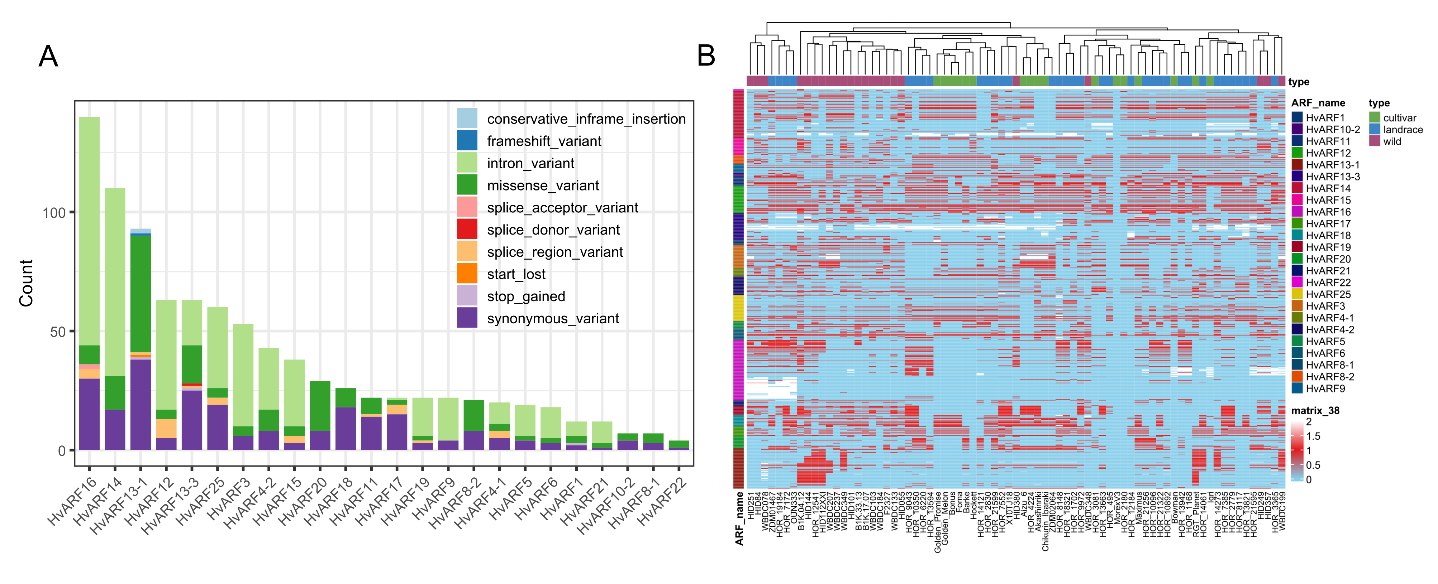

**Supplementary figure 4: Re-mapping and variation detection of *ARF* genes across 76 barley genotypes based on the MorexV2 reference genome.** (A) Number of SNPs annotated within each *ARF* gene across the pan-genome panel. (B) SNP matrix displaying genotypic variation across 76 barley accessions. Red indicates the alternative allele, blue represents the reference allele, and white denotes heterozygous calls.

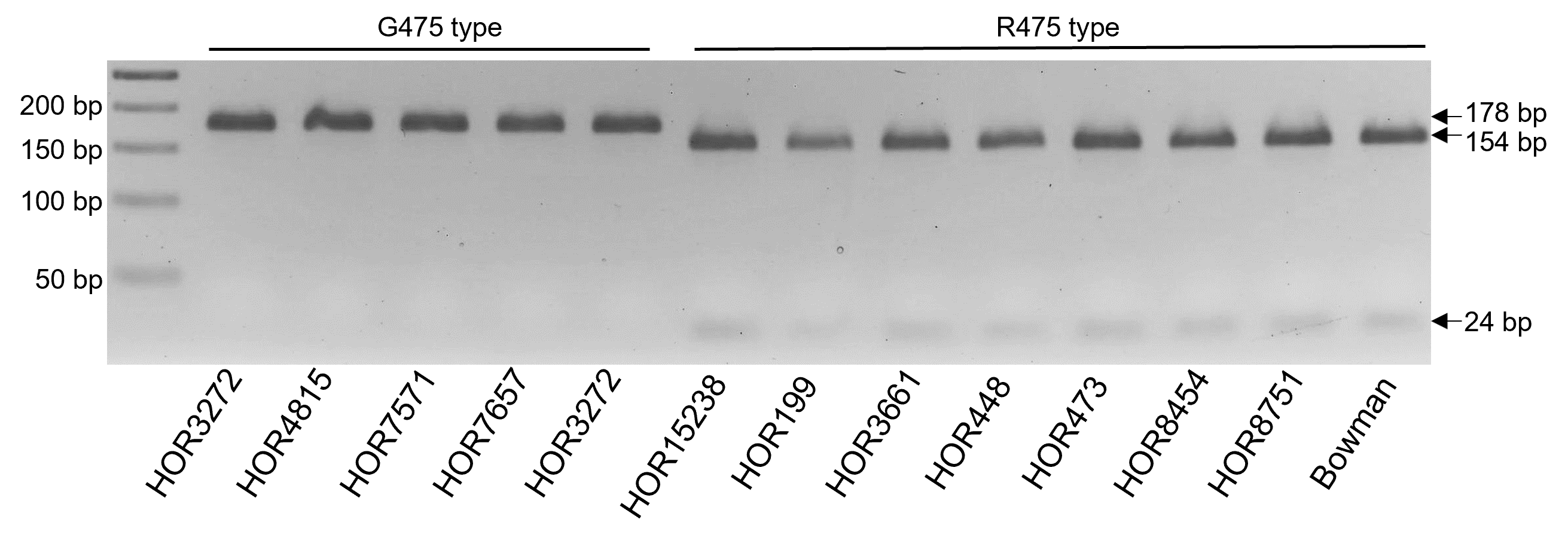

**Supplementary figure 5: Validation of the dCAPS marker for distinguishing HvARF3 alleles at the R475G variant.** PCR products were digested with the restriction enzyme *DdeI*. The G475 allele (non-preferred) remains uncut, resulting in a single 178 bp band, while the R475 allele (preferred) is cleaved into two fragments of 154 bp and 24 bp. Agarose gel electrophoresis of digestion products from representative barley genotypes confirms the efficacy of the marker in distinguishing allelic variants.
